# Lysosome exocytosis is required for mitosis

**DOI:** 10.1101/375816

**Authors:** Charlotte Nugues, Nordine Helassa, Dayani Rajamanoharan, Robert D Burgoyne, Lee P Haynes

## Abstract

Mitosis, the accurate segregation of duplicated genetic material into what will become two new daughter cells, is accompanied by extensive membrane remodelling and membrane trafficking activities. Early in mitosis, adherent cells partially detach from the substratum, round up and their surface area decreases. This likely results from an endocytic uptake of plasma membrane material. As cells enter cytokinesis they re-adhere, flatten and exhibit an associated increase in surface area. The identity of the membrane donor for this phase of mitosis remains unclear. Here we show by biochemical and imaging approaches that lysosomes undergo exocytosis at telophase and that this requires the activity of phosphatidylinositol 4-kinase-IIIβ. Inhibition of lysosome exocytosis resulted in mitotic failure in a significant proportion of cells suggesting that this facet of lysosome physiology is essential and represents a new regulatory mechanism in mitosis.

## Introduction

The emergence of complex multicellular organisms including humans has rested upon the evolution of mechanisms by which single cells can duplicate themselves to build distinct organs and tissues. The process of mitosis in higher organisms is responsible for normal growth, development, aging and repair of most body structures and defects during this event can lead to disease states including cancer (Dominguez-Brauer *et al.*, 2015). Understanding how mitosis is controlled is therefore fundamental to tackling a number of prevalent human developmental disorders and diseases.

It has long been known that mitosis involves dramatic alterations in cellular morphology and, particularly for adherent cell types, there is a weakening in attachment to the extracellular matrix on entry in to mitosis. This is accompanied by a significant reduction in plasma membrane (PM) surface area that manifests as cell rounding. These morphological changes appear intimately linked to activation of specific cell signalling pathways and cytoskeletal rearrangements occurring within the cytoplasm (Meyers *et al.*, 2006). It was originally speculated that changes in cell volume and surface area could be achieved through folding and unfolding of the PM perhaps into villus like protrusions/invaginations and some cell types may employ this strategy (Porter *et al.*, 1973; Erickson and Trinkaus, 1976). Intuitively it was speculated that an alternative mechanism for dynamic adjustment of PM surface area during mitosis could occur through the processes of endo- and exocytosis (Erickson and Trinkaus, 1976). The first studies examining vesicular trafficking during mitosis identified endocytic compartments as potential sources of membrane for surface area expansion late in mitosis (Boucrot and Kirchhausen, 2007) while later work identified Golgi-derived vesicles as possible mediators of this phenomenon (Goss and Toomre, 2008).

Lysosomes are remarkably versatile intracellular organelles that perform numerous essential functions within cells (Luzio *et al.*, 2007). They are master regulators of cellular metabolism, contribute to catabolism of cellular debris, including foreign pathogens, and are central to macro-autophagy (Wartosch *et al.*, 2015). In addition lysosomes are important calcium (Ca^2+^) signalling platforms (Kilpatrick *et al.*, 2013) and can facilitate repair of mechanical damage to the PM in most cell types (Rodriguez *et al.*, 1997). This latter function of lysosomes depends upon their ability to undergo Ca^2+^-dependent exocytosis at the PM and a combination of lysosomal membrane material and soluble lysosomal hydrolases appear important in the process of wound healing (Castro-Gomes *et al.*, 2016). More recently it has been proposed that excessive lysosomal exocytosis may in-part be responsible for the enhanced invasiveness of cancer cells (Liu *et al.*, 2012; Machado *et al.*, 2015). In work examining the role of a small Ca2+ binding protein, CaBP7 and phosphatidylinositol 4-kinaseIIIβ (PI4K) our laboratory uncovered a potential but undefined role for lysosomes during mitosis in mammalian cells (Rajamanoharan *et al.*, 2015). We characterised a highly organised and temporally coordinated clustering of lysosomes to the site of PM constriction in mitotic cells prior to cytokinesis. Inhibition of this clustering, through a PI4K-dependent mechanism, was accompanied by a significant increase in mitosis failure. Taken together these observations indicated that the spatial distribution of lysosomes had an important functional impact on mitosis progression. However, the exact role of lysosomes during mammalian the steps leading to cytokinesis remains unclear. In the present study, with the knowledge that mitosis requires membrane remodelling and that an organelle(s) with the ability to undergo exocytosis could be important for PM expansion we have examined whether lysosomes undergo exocytosis during mitosis and whether this is an essential requirement. Using a number of mammalian cell-lines we show, using biochemical assays that track the release of soluble lysosomal content into the extracellular space, that there is an increase in lysosome exocytosis as cells undergo mitosis. These data were confirmed in Total Internal Reflection Fluorescence Microscopy (TIRFM) imaging studies where we monitored coordinated lysosome (organelles identified based on their ability to accumulate acidotropic dye but which were negative for mannose-6-phosphate receptor) trafficking and fusion events at the site of PM constriction. Pharmacological inhibitors targeting the exocytic machinery or PI4K directly induced significant mitosis failure suggesting that lysosome exocytosis is functionally required for normal mitosis completion. These findings provide new insights into the cell biology of mitosis and it is possible that this behaviour of lysosomes could represent a new way to target rapidly proliferating cells to induce mitotic arrest.

## Results

### Lysosomal protein expression and surface appearance is upregulated in mitotic cells which exhibit a corresponding increase in lysosomal exocytosis

We first examined if there were discernible differences in lysosomal protein expression and cellular distribution between asynchronous and chemically synchronised (thymidine-nocodazole) cells (Figure 1A). Surface proteins were labelled with a cell-impermeant biotinylated cross-linking reagent (Sulfo-NHS-SS-biotin), streptavidin affinity-purified from detergent extracted lysates and analysed by Western blotting for the lysosomal membrane protein LAMP-1 (Figure 1A). Densitometry analysis of this data highlighted that intracellular LAMP-1 protein levels in mitotic cells (released from the nocodazole block for 45 minutes, typically cells have reached late anaphase/early telophase by this time) increase 6.7-fold compared to levels in asynchronous cells. (Tubulin loading control normalised, Figure 1A). In mitotic cells, surface LAMP-1 represented 8% of the total LAMP-1 signal (surface + cytosol) in comparison to 0.28% in asynchronous cells.

**Figure 1.**
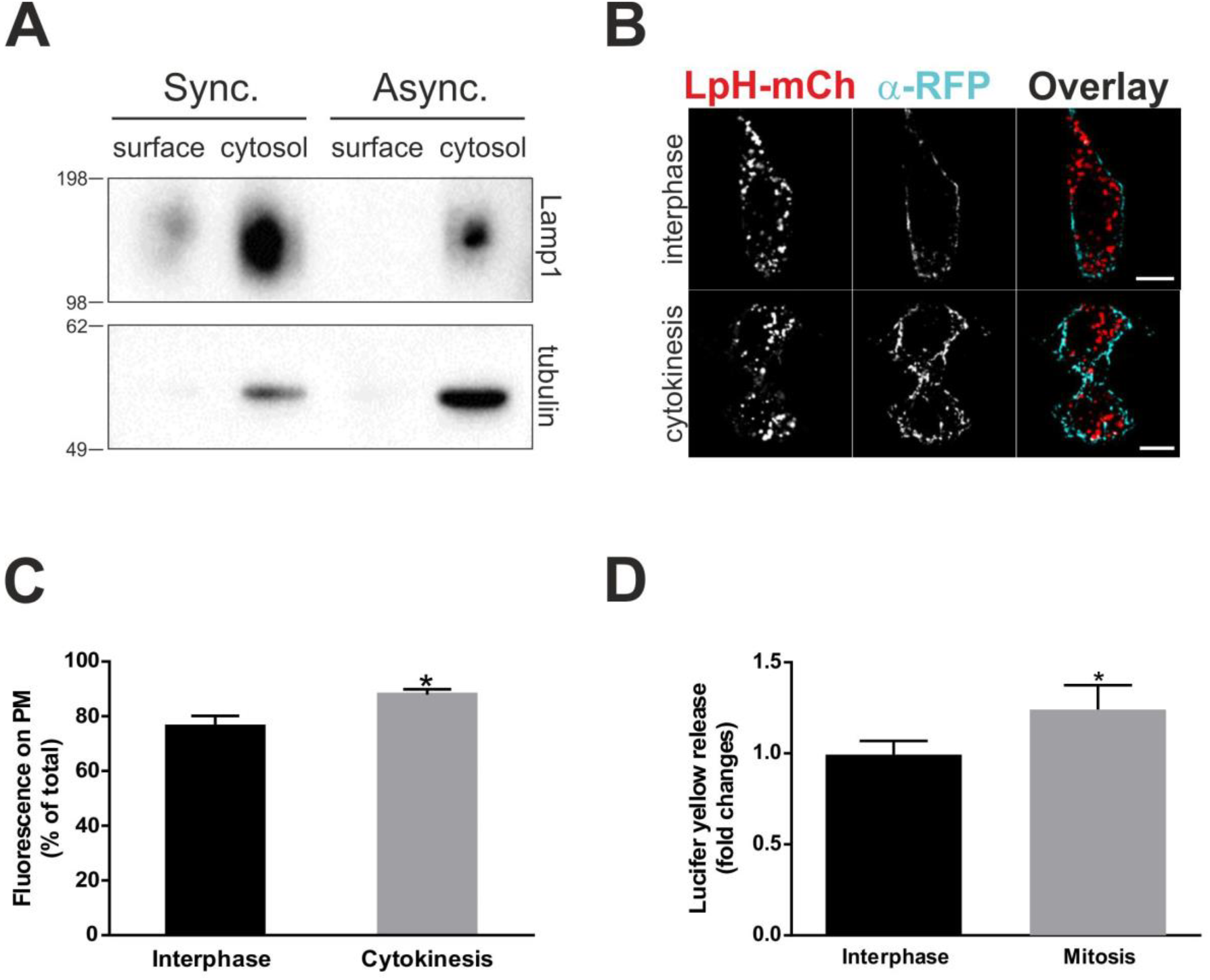
Cellular expression and plasma membrane residency of a lysosomal membrane protein increases during mitosis. A) Western blot analysis of surface and cytosol fractions revealed increased expression of LAMP-1 in the cytosol of mitotic cells and its appearance at the surface of synchronised HeLa cells but not at the surface of interphase cells. The surface accessible protein fraction of synchronous (sync) and asynchronous (async) cells was captured by surface biotinylation and streptavidin affinity purification. Post-strepavidin cell lysates (non-surface, cytosol) were also prepared. Cytosolic LAMP-1 signals were normalised to a β-tubulin loading control signal from the same samples. Predicted sizes: 120 kDa (LAMP1), 50 kDa (β-tubulin). B) Confocal images show deconvoluted middle sections at interphase and cytokinesis. HeLa cells were transfected with LpH-mCh. The luminal part of the construct (mCherry, red) was detected on the cell surface using rabbit anti-RFP (blue) and goat anti-rabbit Alexa 405 conjugated antibodies, prior to cell fixation without cell permeabilisation. Scale bars: 10 μm. C) Quantification of data from (B). The fluorescence of RFP at the membrane expressed as % of total cellular RFP fluorescence is increased in the late stages of cell division, compared to HeLa cells in interphase. *p = 0.0446; n = 5 cells in interphase; n = 4 cells in late telophase/cytokinesis. D) *Lucifer yellow (LY) secretion increases during mitosis in BSC-1 cells*. All cells were synchronised using a double thymidine block and loaded with 1mg/ml LY (12-hour pulse and 2-hour chase). After the second thymidine treatment, cells were maintained in thymidine (interphase) or released from the block (mitosis). LY secretion was assessed immediately after release (time = 0 hours) and 20 hours post-release in both mitotic and interphase cell populations. Data were corrected for cell death using bright field live cell images (Fig. extended view 2). The results are depicted as LY fold changes, from 0 to 20 hours after release. Results shown as mean ± S.E.M. Unpaired student t-test *p = 0.0494, N = 3.

As a complimentary method to assess lysosomal protein at the PM we used surface immunofluorescence. Due to the difficulty of finding reliable antibodies directed towards luminal epitopes of lysosomal proteins, we developed LysopHluorin-mCherry (LpH-mCh, Figure S1A), an mCherry tagged, lysosomally targeted construct derived from the previously described Lyso-pHoenix (Rost *et al.*, 2015). LpH-mCh contains the CD63 lysosome targeting information followed by pHluorin (a pH sensitive ecliptic GFP variant quenched at acidic pH) and mCherry (a pH insensitive red fluorophore, pKa ~4.5) such that both tags are located within the lysosome lumen. LpH-mCh targets extensively to structures positive for the acidotropic dye Lysotracker® but largely negative for mannose-6-phosphate receptor (Figure S1B and S1C) indicating that LpH-mCh is a lysosome-specific pH sensor. In cells expressing LpH-mCh, if a lysosome were to exocytose at the PM then the pHluorin and mCherry tags would become accessible at the cell surface to specific antibodies. We used mCherry fluorescence to determine LpH-mCh localisation throughout the cell and anti-RFP antibody for staining of non-permeabilised cells to detect surface-accessible mCherry protein (Figure 1B). We normalised the surface LpH-mCh signal to total cellular LpH-mCh signal (PM + intracellular) and observed a statistically significant increase in surface LpH-mCh signal in cells at cytokinesis (Figure 1C), consistent with our surface biotinylation data (Figure 1A).

Having biochemical and imaging-based evidence that lysosomal membrane protein abundance at the PM apparently increased in mitotic cells we next wanted to test if this was due to lysosome exocytosis and not biosynthetic trafficking of newly synthesised LAMP-1 to lysosomes. The latter interpretation would be unusual since the Golgi apparatus fragments early during mitosis (Acharya *et al.*, 1998) precluding synthesis of new secretory pathway proteins and therefore would require a novel mechanistic explanation. We measured lysosome exocytosis directly using a previously characterised Lucifer Yellow (LY) uptake and secretion assay (Rodriguez *et al.*, 1997). For these experiments we used BSC-1 cells (African green monkey kidney epithelial cells) since HeLa cells have previously been shown to exhibit relatively low levels of lysosome exocytosis (Rodriguez *et al.*, 1997). LY exocytosis was determined at different time points: immediately (directly following release of the second thymidine block in mitotic cells (t = 0 hours)) or at 20 hours following the release (t = 20 hours), to permit synchronised cells to undergo a complete round of division. Cells arrested in interphase were processed at precisely the same time points however they were maintained in the presence of thymidine for the entire experiment. Mitotis- and interphase-specific LY release was then calculated as fold changes between the t = 0 hour and t = 20hour time points (Figure 1D). Cells demonstrated a significant increase in exocytosis during mitosis, from a 0.99-fold increase in interphase to a 1.24 fold at mitosis (Figure 1D). All data were corrected for cell death by examination of cell density in bright-field images.

### Lysosome exocytosis visualised by TIRFM in mitotic cells

Having acquired data indicating that lysosome exocytosis increases as cells progress through mitosis we examined this process at the level of individual exocytic events in live cells using TIRFM. In mitotic cells expressing LpH-mCh or loaded with Lysotracker® we observed a remarkable and highly dynamic clustering of lysosomes first to the site of PM constriction early in telophase and subsequently to either side of the intercellular bridge (as previously reported using standard confocal microscopy (Rajamanoharan *et al.*, 2015)) (Figure 2A, B and Movies S1, S2 and S3). Employing diffusion analysis developed in (Goss and Toomre, 2008) (see methods) to determine vesicle-PM fusion (exocytosis) we demonstrated that lysosomes underwent fusion events consistent with exocytosis at the PM in these cells (Figure 2C and D and Movies S4 and S5). Over similar time courses in interphase cells we were unable to observe individual lysosome exocytosis events. Examination of TIRFM data showed that we were however able to detect numerous docking/undocking like events (movement of lysosomes towards and away from the PM) that were not accompanied by exocytosis in both mitotic and interphase cells (Figure 2E and movie S6). In conjunction with diffusion analysis, we used the complimentary measurement of pHluorin de-quenching as a readout of lysosome exocytosis at the PM. To validate de-quenching as a diagnostic tool we first characterised exocytosis in cells at interphase, treated with 10 μM ionomycin (Figure S2 and movie S7). Upon fusion of lysosomes with the PM, the organelle lumen and the extracellular space become continuous, resulting in lysosomal alkalinisation and a marked increase in pHluorin fluorescence (Figure 2C and movies S4 and S5). Collectively these data show that lysosomes are highly mobile at all stages of the cell cycle, cluster at either side of the intercellular bridge at telophase and exocytose during mitosis.

**Figure 2.**
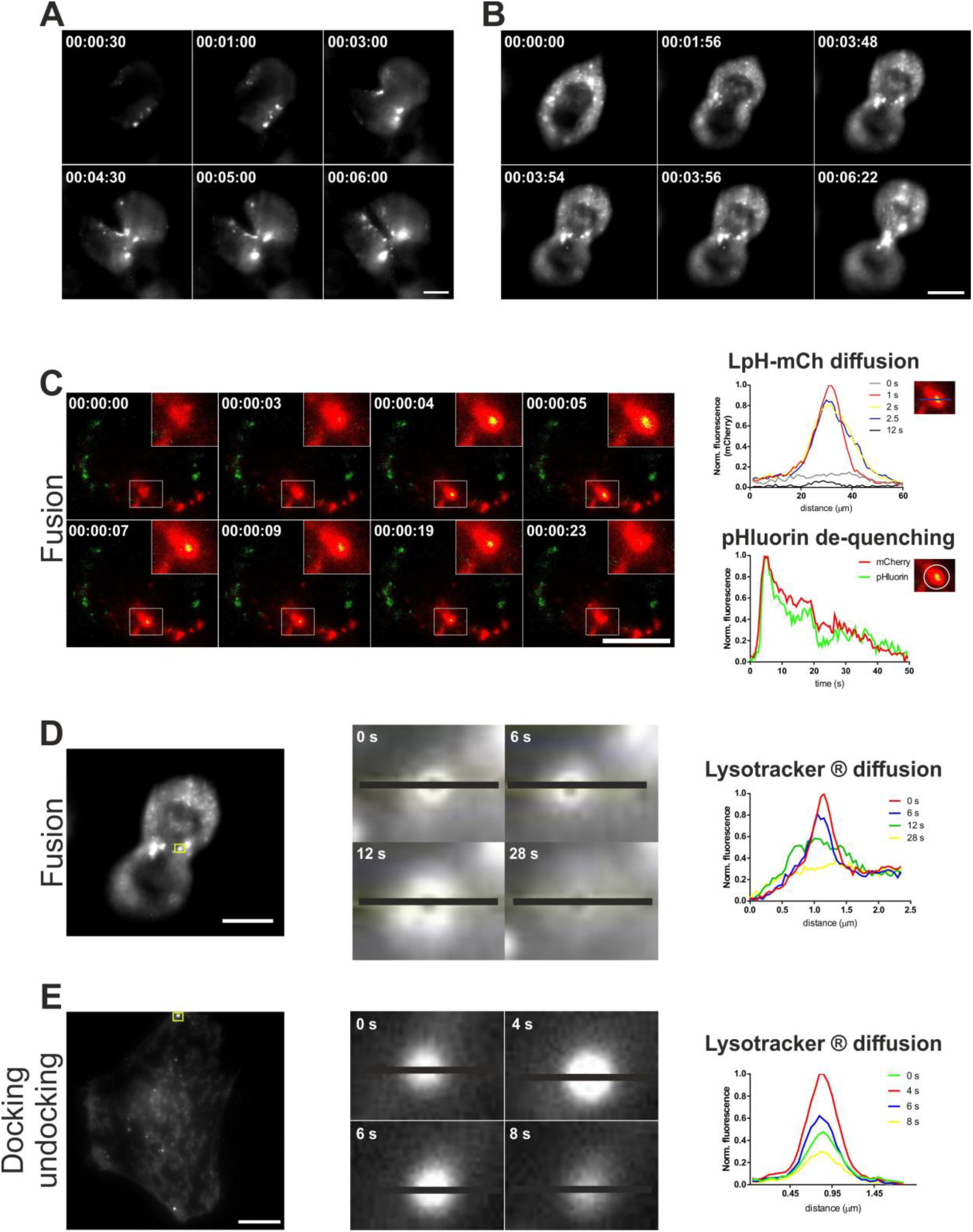
TIRFM analysis of lysosomes located near the plasma membrane which localise to the site of cytoplasmic constriction and undergo exocytosis during mitosis. A & B) Cells were loaded with 50 nM Lysotracker®-red for 30 minutes and imaged at a rate of 1 frame every 30 seconds (BSC-1 cells; A) or every 2 seconds (NRK cells; B) showing the clustering of lysosomes during mitosis. C) Example of lysosome exocytosis in a LpH-mCh-stable NRK cell undergoing cytoplasmic constriction (late telophase/early cytokinesis). The white square (and zoomed view) represents the exocytosing lysosome analysed in the accompanying graphs. The top graph shows the diffusion analysis along a line (black) drawn across the lysosome. The second graph illustrates the sharp and simultaneous increase in both pHluorin and mCherry fluorescence upon exocytosis in a region of interest drawn around the same lysosome. LpH-mCh-NRK cell was imaged every 0.5s. D) Zoomed view of a lysosome loaded with Lysotracker® (highlighted by the yellow square). The time indicates the time for the lysosome to dock, fuse with the PM and diffuse Lysotracker® fluorescence in the extracellular space. The graph depicts the spreading of the fluorescence across the black line. E) Diffusion analysis of a Lysotracker®-loaded lysosome moving in close proximity to cytoplasmic face of the PM (docking-undocking) during interphase. NRK cell was imaged every 2 s. The lysosome analysed is highlighted by the yellow square and represented in the zoomed view. The graph demonstrates fluorescence does not spread across the line drawn across the lysosome (blue line) as the width of fluorescence remains unchanged. The time specifies the duration of the lysosome vertical movement, which is shorter than fusion duration. Time indicated as hr:min:sec. White scale bars: 10 μm.

### Inhibition of lysosome exocytosis with N-ethylmaleimide or PIK-93 induces mitotic failure

Mitosis represents a small fraction of the total cell cycle and typically takes cultured cells no more than one hour to complete. To probe the functional requirement of lysosome exocytosis specifically during the small time window encompassed by mitosis we decided to employ acute treatment of cells with pharmacological inhibitors. Silencing of lysosomal SNARE proteins to inhibit exocytosis was considered but rejected as an experimental approach for two reasons: 1) Lysosomal SNAREs such as VAMP7 are also known to be abundant on late endosomes (Chiaruttini *et al.*, 2016). Therefore, siRNA depletion of this protein would potentially influence the behaviour of endocytic compartments other than lysosomes; 2) siRNA would also require transfection of cells for many days which would increase the chances of observing effects on lysosome physiology unrelated to mitosis.

N-ethylmaleimide is a well-documented inhibitor of vesicular fusion events dependent upon the hexameric ATPase NSF (Block *et al.*, 1988; Sivaramakrishnan *et al.*, 2012). We acutely treated LY loaded cells with 1mM NEM for 15 minutes to minimise effects on other cellular processes and assessed the impact on lysosome exocytosis and mitosis (Figure 3A & 3B). NEM treatment reduced lysosome exocytosis by approximately 40% (Figure 3A) and this correlated with a complete block on cell division monitored in live cell analysis of NEM treated mitotic cells (Figure 3B). NEM is known to disrupt multiple NSF dependent trafficking events and as such is a non-specific inhibitor. Based on previous data generated in our laboratory suggesting a role for PI4K in lysosome physiology, particularly in lysosome clustering prior to cytokinesis and progress of mitosis, we tested the effect of the PI4K inhibitor PIK-93 (Rutaganira *et al.*, 2016), on lysosome exocytosis (Figure 3C). Although PIK-93 may not be completely specific for PI4K it is considerably more selective than NEM and was used at the published IC50 for PI4K (19 nM) which is significantly below the inhibitory concentration for other kinases. PIK-93 treatment of cells abolished Ca^2+^-dependent lysosome exocytosis of the soluble lysosomal hydrolase N-acetyl β-D Glucosaminidase (NAG)(Rodriguez *et al.*, 1997) (Figure 3C) and significantly increased the proportion of polynucleate cells (Figure 3F). Having shown that inhibition of PI4K impairs exocytosis, we next examined the effect of PI4K overexpression on lysosomal fusion with the PM. Although detection of changes in lysosome exocytosis remained challenging in HeLa cells, overexpression of PI4K triggered a surprisingly significant increase in NAG release, further reinforcing the involvement of the enzyme in the exocytic process (Figure 3D).

**Figure 3.**
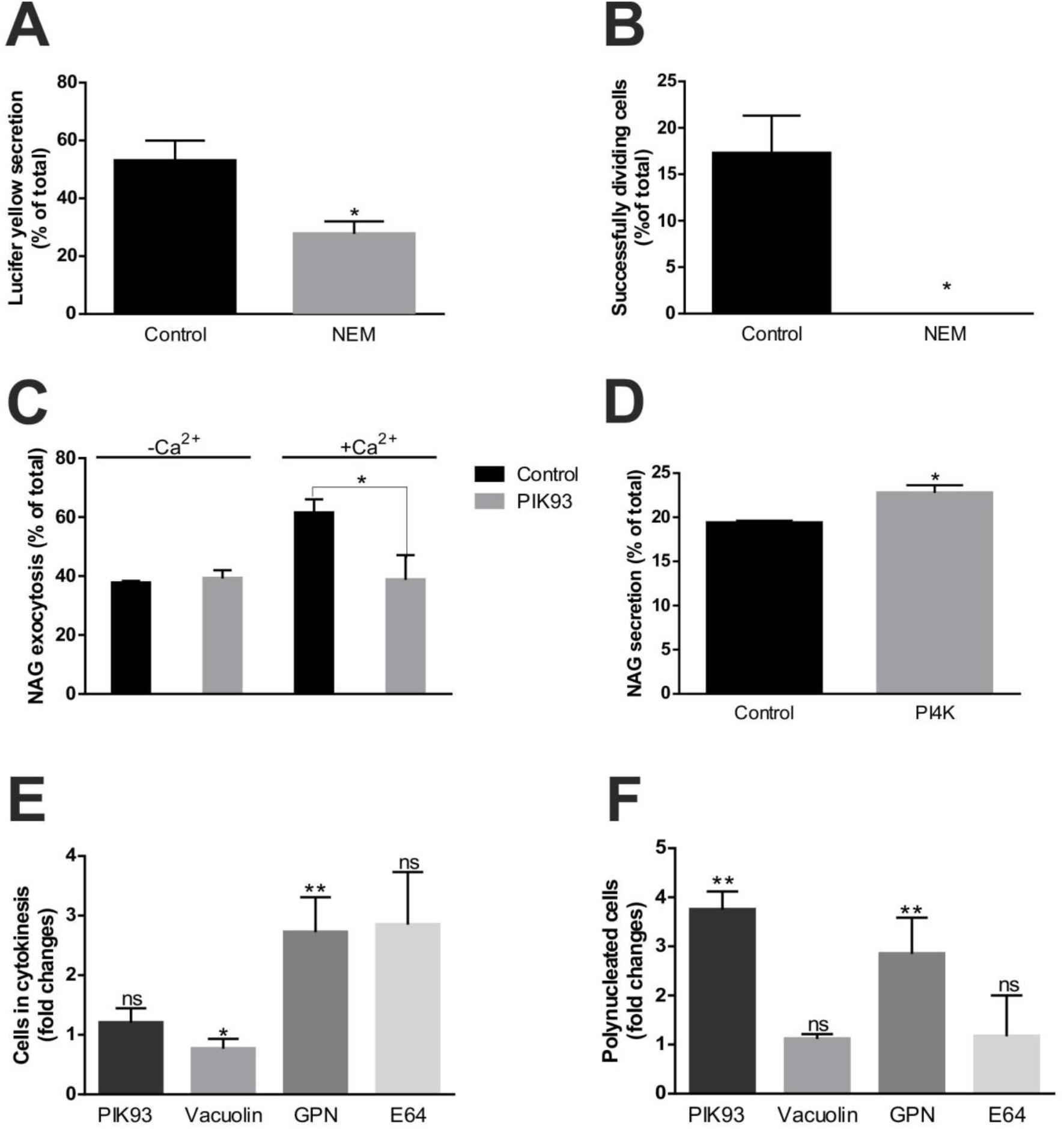
Lysosome exocytosis requires PI4K and is essential for mitosis completion in mammalian cells. A) NEM reduces LY release in BSC-1 cells. Cells were incubated with 1mg/ml LY for 4 hours, washed and chased for a further 2 hours to permit accumulation in lysosomes. Cells were subsequently treated with 1 mM NEM (or 0.25% (v/v) Ethanol, vehicle control) for 15 min. LY secretions were then measured and expressed as % of total cellular LY; *p = 0.0370, N = 3. B) NEM treatment blocks mitosis in BSC-1 cells. Cells were treated with 1 mM NEM (or 0.25% (v/v) Ethanol, vehicle control) for 15 min and imaged using a 3i spinning disk microscope. The proportion of cells reaching mitosis completion was counted and expressed as % of total cell numbers in each field of view. *p = 0.0130, n(control) = 247 cells, n(NEM) = 217 cells, N = 3. C) PI4K inhibition by PIK93 decreases Ca^2+^-dependent exocytosis in NRK cells. Cells were incubated with 19 nM PIK93 (or 0.1% (v/v) DMSO, control) for 40 minutes in the presence or absence of 10 M ionomycin for 10 min. NAG lysosomal enzyme secretion was measured in the incubation media and in the cell lysates and calculated as % of total cellular NAG. *p = 0.0149, N = 3. D) Overexpression of PI4K enhances lysosome exocytosis. Lysosomal NAG release was assessed in HeLa cells transfected with HA-PI4K (control: transfection reagent only). *p = 0.0356, N = 3. E) and F) Cells were treated for 40 min with 19 nM PIK93 (or 0.1% (v/v) DMSO, control) or overnight with 1 μM Vacuolin (or 0.1 % DMSO, control), 50 μM GPN (or 0.1% (v/v) DMF, control) or 100 μM E64 (or untreated, control). Cells were then fixed, permeabilised and immunostained with anti-tubulin antibody and DAPI. The proportion of cells at cytokinesis (E) and bi/tetranucleate cells (F) were counted in each condition. Results represent fold changes under treatment with the drugs compared to their respective vehicle controls. The proportion of bi/tetranucleate NRK cells is significantly raised under treatment with PIK93 (**p = 0.0031, n(control) = 942 cells, n(PIK93) = 1310 cells, N = 6). No significant mitosis defects were observed in NRK cells treated with the lysosomal cysteine protease inhibitor E64 (n(control) = 521 cells, n(E64) = 619 cells, N = 6). GPN but not Vacuolin triggered significant mitosis defects in HeLa cells (*p(vacuolin) = 0.0298, **p(GPN, cytokinesis) = 0.006, **p(GPN, polynucleated) = 0.001, n(vacuolin) = 1560 cells (n(control) = 1250 cells) n(GPN) = 1648 cells (n(control) = 1322 cells), N(cytokinesis) = 12, N(polynucleated) = 8. Statistical significance was calculated using unpaired student t-test and all results are shown as mean ± S.E.M. ns: not significant.

In order to test whether lysosomal hydrolases released during exocytosis are required mitosis completion we treated cells with the cathepsin inhibitor E64 (Figure 3E, 3F and Figure S3). These cells exhibited no detectable impairment in mitosis compared with controls. Glycyl-L-phenylalanine 2-naphthylamide (GPN) is another reagent that has been used to interfere with lysosome function (Jadot *et al.*, 1984; Berg *et al.*, 1994). It is a specific cathepsin C substrate which, once cleaved, is osmotically active within the lysosome lumen leading to excessive water accumulation and physical swelling/perforation of the organelle. Cells were treated overnight with GPN and then fixed for immunofluorescence analysis with an anti-tubulin antibody and DAPI staining. In a previous report we described an analysis of mitotic failure based on the appearance of cells with 2n copies of the nucleus or an increased frequency of cells connected by an intercellular bridge (Rajamanoharan *et al.*, 2015). We used the same approach here to examine the effect of inhibiting lysosome activity with GPN. GPN treatment induced a significant increase in the proportion of bi-/tetra-nucleate cells and cells connected by a clearly identifiable intercellular bridge (Figure 3E and 3F). Another pharmacological agent suggested to impair lysosome function, vacuolin-1, had no effect on mitosis (Figure 3E and 3F) which is consistent with a previous report showing that this drug does not inhibit lysosome exocytosis (Huynh and Andrews, 2005).

## Discussion

Understanding the regulatory mechanisms that govern mitosis has, as well as revealing new cell biological insights, potential impact for being able to address human health issues. The Spindle Assemble Checkpoint (SAC) is recognised as the most important regulatory mechanism that operates during mitosis (Rieder and Maiato, 2004; Lara-Gonzalez *et al.*, 2012). SAC is a valid anti-cancer target and a number of currently employed chemotherapeutic agents work by stabilising or destabilising the microtubule network (MT) to activate this checkpoint and stall cells undergoing division (Gascoigne and Taylor, 2009; Dominguez-Brauer *et al.*, 2015). Blocking mitosis in this way has the immediate effect of halting cell proliferation but has an added advantage in that such cells can enter mitotic cell death ultimately destroying transformed cells. Drugs such as Taxol (MT stabiliser) and the vinca alkaloids (MT destabilsers) have been employed clinically for many decades however they have potential off-target effects on normal cells and therefore new generation anti-mitosis agents will ideally exhibit greater cancer cell selectivity (Dominguez-Brauer *et al.*, 2015). One way in which specificity can be enhanced is to identify new regulatory inputs required for successful mitosis completion.

Roles for phosphoinositide metabolism (Brill *et al.*, 2011; Echard, 2012), calcium signalling (Whitaker, 2006) and vesicular membrane trafficking (Boucrot and Kirchhausen, 2007; Goss and Toomre, 2008) during mitosis have been reported although they are significantly less well understood. Control of membrane remodelling during mitosis is accepted as integral for normal cell division and manipulations that prevent cell-rounding place a block on mitosis (Boucrot and Kirchhausen, 2007). The underlying cell biology mediating membrane contraction and expansion have been investigated in previous studies and it has been suggested that multiple membrane trafficking pathways involving Golgi vesicles or endosomes could be responsible (Boucrot and Kirchhausen, 2007; Goss and Toomre, 2008). These reports provided unique insights into organelle exocytosis during mitosis but they did not prove unequivocally that either Golgi vesicles or endosomes are essential for normal mitosis.

In a previous study, we reported a distinct redistribution of lysosomes in mitotic HeLa cells which appeared to have functional relevance for mitosis completion albeit in an undefined manner (Rajamanoharan *et al.*, 2015). Exocytosis of lysosome related organelles is important in a number of professional secretory cell types, particularly those of the immune system (Marks *et al.*, 2013). In pioneering studies the work of the Andrews laboratory described a more general feature of standard lysosomes, present in almost all cell types, to undergo Ca2+- dependent exocytosis in response to PM damage (Rodriguez *et al.*, 1997; Reddy *et al.*, 2001). These observations uncovered a previously unknown and unexpected property of lysosomes, which has since been implicated in cancer cell migration and drug resistance (Liu *et al.*, 2012; Machado *et al.*, 2015; Zhitomirsky and Assaraf, 2017). It is clear that lysosomes serve multiple important functions in cells with increasing attention being paid to their influence on cancer cell behaviour.

In this study we describe how a known lysosome function acts in a hitherto unknown but essential area of mammalian cell biology. We have characterised lysosome exocytosis in various mammalian cell types undergoing mitosis and show that disruption of this function impairs normal mitosis completion. Our data suggest that lysosomes serve a key function during mitosis that requires their exocytosis at the cell PM. One interpretation of our results is that, under normal conditions, lysosome exocytosis is linked to the PM remodelling, particularly membrane expansion, that occurs during telophase in mitotic cells. This is the first study to combine biochemical analyses of organelle exocytosis with high resolution imaging to describe how lysosomes might contribute additional membrane during mitosis. Lysosome exocytosis as a general mechanism for PM wound repair in various cell types was characterised over twenty years ago (Rodriguez *et al.*, 1997). The prevailing model posited that lysosomal membrane was incorporated into the lesion to restore PM integrity (Reddy *et al.*, 2001). This view has been refined by more recent reports showing that soluble lysosomal hydrolases released upon lysosome exocytosis are key extracellular factors in PM wound repair (Tam *et al.*, 2010; Castro-Gomes *et al.*, 2016). Acid sphingomyelinase (Tam *et al.*, 2010) and the cysteine proteases cathepsin B and L (Castro-Gomes *et al.*, 2016) have been shown to be required for PM repair. We examined whether the same machinery that is involved in PM wound repair also operates during mitosis by treating cells with the cathepsin inhibitor E64 (inhibitor of cysteine active site cathepsins including B, L and K isoforms (Matsumoto *et al.*, 1999)) and assessing the impact on mitosis completion. Since E64 treatment did not inhibit mitosis this suggests that the exocytosis of lysosomes during mitosis serves a mechanistically distinct function from that which operates during PM wound repair which we interpret as membrane donation. Previous work in our laboratory tested the effect of Pepstatin-A (aspartate active site cathepsin inhibitor – isoforms D and E (McGlinchey and Lee, 2015)) on mitosis progression and observed no detectable effect consistent with a dispensable role for lysosomal proteolytic activity during mitosis (data not shown).

Lysosome exocytosis during mitosis is also unique in that it is a PI4K dependent process. PI4K is responsible for generating phosphatidylinositol-4-phosphate, a fusogenic lipid, and has a well-documented activity in the biogenesis of anterograde transport vesicles at the Trans-Golgi Network (Haynes *et al.*, 2005). A previous study characterised a biochemically distinct pool of PI4K on lysosomes (Sridhar *et al.*, 2013) and our observations indicate that a potential function of the kinase on these organelles is to prime them for exocytosis at the PM. The data presented here are consistent with our previous work where we demonstrated that expression of CaBP7, a PI4K inhibitor, disrupts lysosome clustering and inhibits mitosis (Rajamanoharan *et al.*, 2015). Interestingly, the effect of PIK-93 observed in this study was larger in NRK cells compared with HeLa cells which may reflect a heightened sensitivity to the drug of cells exhibiting higher levels of Ca^2+^-stimulated lysosome exocytosis. This suggests that PI4K has a key role in lysosome function during mitosis in mammalian cells and that it is required for Ca^2+^-dependent exocytosis. The role of PI4K in lysosome exocytosis may not be restricted exclusively to mitosis and could be required for exocytosis of these organelles more generally, including during PM wound repair, however, to date, this possibility has yet to be investigated.

Collectively, our data highlight a novel role for lysosome exocytosis in mitosis involving PI4K, a process that represents a potentially new regulatory input controlling mitosis progression, and which may offer an additional means to enhance the specificity in targeting rapidly proliferating cells.

## Material and methods

### Cell culture

HeLa, HEK293T and NRK-49F cells were maintained in HN media (DMEM supplemented with 10% (v/v) Foetal Bovine Serum (FBS), 1% (v/v) non-essential amino acids (NEAAs) and 1% (v/v) penicillin/streptomycin (P/S). BSC-1 cells were obtained from ATCC and maintained in EMEM supplemented with 10% (v/v) FBS, 1% (v/v) NEAAs and 1% (v/v) P/S. All cells were cultured at 37°C, 95% (v/v) air/5% (v/v) CO_2_.

### Cloning of LpH-mCh

LysopHluorin-mCherry (LpH-mCh) was generated by PCR amplification of the CD63-pHluorin domain of pCMV-lyso-pHoenix (from Addgene clone 70112; forward primer (NheI): 5’- CT<u>GCTAGC</u>ATGGCGGTGGAAGGAGGAATG-3’; reverse primer (BamHI): 5’- CT<u>GGATCC</u>TTTTTGTATAGTTCATCCATGCCATG-3’). The fragment was digested with BamHI HF and NheI HF (NEB) and ligated into the pmCherry-N1 vector (Clontech) using T4 DNA ligase (NEB). Recombinant plasmids were verified by DNA sequencing (Source Bioscience, UK). Correct expression of LpH-mCh in HeLa cells was verified by Western blot and immunofluorescence analysis.

### Generation of pHIV-LpH-mCh and a LpH-mCh stable NRK cell line using lentivirus

pHIV-dTomato (Addgene) was digested with XmaI and ClaI (NEB) to excise the dTomato fragment. The LpH-mCh coding sequence was amplified by PCR using primers designed for In-Fusion® HD Cloning, so that the resulting fragment shared 15-bp homology with the pHIV vector (forward primer: 5’-TTCTAGAGTACCCGGATGGCGGTGGAAGGAGGAATG-3’; reverse primer: 5’-AGGTCGACGGTATCGCTTGTACAGCTCGTCCATGCC-3’).

XmaI/ClaI Digested pHIV-dTomato and LpH-mCh PCR product were combined using In-Fusion® HD enzyme (100ng linearised pHIV, 1μl PCR product, 2μl enzyme; reaction allowed for 15 min at 50°C). Recombinant LpH-mCh plasmid was amplified by standard techniques and verified by DNA sequencing (Source Bioscience, UK).

HEK293T cells were grown in 10 cm cell culture dishes to 80% confluence and transfected with lentiviral components and pHIV-lpH-mCh using Lipofectamine® 2000 (Thermo Fisher Scientific) according to manufacturer’s instructions. pMDLg/pRRE was a gift from Didier Trono (Addgene plasmid # 12251), pRSV-Rev was a gift from Didier Trono (Addgene plasmid # 12253) and pCMV-VSV-G was a gift from Bob Weinberg (Addgene plasmid # 8454). Briefly, 20 μg pHIV-lpH-mCh, 10 μg pMDLg/pRRE, 5 μg pRSV-Rev and 6 μg pCMV-VSV-G were added to 100 μl Lipofectamine® 2000 in Opti-MEMTM (ThermoFisher Scientific). The solution was incubated for 20 min at room temperature and 2 ml was added dropwise onto HEK293T cell medium. The virus-containing medium was collected 48 hours post-transfection and centrifuged for 15 min at 1,000 rpm. The supernatant was filtered through a 0.45 μm filter membrane and mixed with fresh medium at a 1:1 ratio. The virus solution was used to infect NRK cells to generate a stable cell line expressing LpH-mCh. Expression of LpH-mCh was verified by confocal imaging and Western blotting.

### Cell Synchronisation

Cells were first arrested at interphase with 2 mM thymidine for a minimum of 20 hours, released from their block with 25 μM deoxycytidine for 3 to 6 hours and blocked again by 2 mM thymidine or 35 ng/ml nocodazole (pro-metaphase block) for 12 to 24 hours. Subsequently cells were either maintained in their block or released with 25 μM deoxycytidine. Prior to all releases, cells were washed 5 times with 10 ml fresh culture medium.

### Biotinylation of cell surface proteins

Synchronous (thymidine-nocodazole treated) and asynchronous HeLa cells were grown to full confluence on 10 cm circular cell culture dishes. Synchronised cells were washed five times with 10 ml HN media to release them from the block and incubated for 1 hr at 37oC in a tissue culture incubator. The culture media was removed from both populations and cells washed with 10 ml of ice-cold phosphate buffered saline (PBS). Cells were gently detached using a cell scraper, transferred to a 15 ml sterile falcon tube and pelleted by centrifugation. Cell pellets were gently resuspended in 1 ml ice-cold PBS supplemented with 0.5 mg/ml EZ-link Sulfo-NHS-SS-biotin (ThermoFisher Scientific) and incubated on a rotator at 4oC for 30 min. Cells were collected by centrifugation and washed once with 1ml ice-cold PBS. Excess unreacted Sulfo-NHS-SS-biotin was quenched by incubation of cells with 1 ml 50 mM NH4Cl in PBS for 30 min on ice. Cells were pelleted by centrifugation (1 min/1000 rpm in a refrigerated microfuge cooled to 4oC) and washed three times with 1 ml ice-cold PBS. Cells were lysed in 1 ml ice-cold PBSL (PBS supplemented with: 0.1% (w/v) sodium dodecyl sulfate (SDS), 1% (v/v) NP-40 and 1% (v/v) Proteoloc protease inhibitor cocktail (Expedeon)) and incubation on a rotator at 4oC for 1 hr. Samples were centrifuged at 13,000rpm, 4oC for 5 min and the supernatants applied to 100μl of Ultralink streptavidin agarose resin (ThermoFisher Scientific) that had been equilibrated by washing three times with 1 ml PBSL. Samples were incubated on a rotator at 4°C for 1hr, streptavidin resin collected by centrifugation (13,000rpm, 4°C, 1 min) unbound material removed to a clean Eppendorf tube (non-surface accessible protein fraction), resin washed five times with 1 ml ice cold PBS and bound proteins extracted by boiling in 100μl dissociation buffer (10% (w/v) sucrose, 10% (v/v) glycerol, 4% (w/v) SDS, 1% (v/v) β-mercaptoethanol, 125 mM HEPES (pH 6.8), 2mM EDTA, 0.05 mg/ml bromophenol blue) surface accessible protein fraction).

### SDS-PAGE and Western blotting

Proteins were resolved on 4-12% Novex NuPage^TM^ Bis-Tris SDS-PAGE and transferred onto nitrocellulose membranes. Filters were incubated with blocking solution (BS: PBS containing 3% (w/v) dried skimmed milk powder) for 1 hour at room temperature. Next, membranes were incubated for 1 hour at room temperature with mouse monoclonal anti-LAMP1 (Santa Cruz; 1:500 dilution) and rabbit anti-β-tubulin (Abcam; 1:1000 dilution) diluted in BS. Membranes were washed 3 times in PBS supplemented with 0.05% (v/v) Tween-20 (PBST), twice with PBS alone and then incubated with goat anti-rabbit and anti-mouse HRP-conjugated secondary antibodies (Sigma, 1:500 dilution in BS) for 1 hour at room temperature. Membranes were then washed three times with PBST and twice with PBS, before addition of equal volumes of enhanced chemiluminescence reagents (Clarity, Biorad). Blots were subsequently developed and imaged using a ChemiDoc-XRS system (BioRad). Densitometry of gel images was performed with ImageJ.

### Immunofluorescence for cytokinesis and polynucleate cell count

HeLa and NRK cells were grown on 13 mm diameter glass coverslips. Cells were washed 3 times in PBS and fixed in 4% (w/v) formaldehyde (Sigma) in PBS for 6 min at room temperature. Cells were washed again in PBS and permeabilised with 0.5% (v/v) Triton-X in PBS for 6 min at room temperature. Following permeabilisation, cells were washed again and blocked for 1 hour at room temperature in 5% (w/v) bovine serum albumin (BSA) in PBS (BSS). Next, cells were incubated in mouse monoclonal anti-tubulin antibody (Abcam DM1A; 1:500 dilution in BSS) for 1 hour at room temperature. Subsequently cells were washed thoroughly in PBS and further incubated with goat anti-mouse Alexa 488 conjugated secondary antibody (1:500 in BSS, Life Technologies) for 1 hour at room temperature. Cells were washed again in PBS, coverslips air dried and mounted onto glass slides using ProLongTM Gold Antifade glycerol containing DAPI (ThermoFisher Scientific).

### Surface staining

Cells were kept on ice at all times and all reagents were maintained at 4°C throughout the whole procedure to inhibit the endocytic process. LpH-mCh-stable NRK cells grown onto 13 mm diameter glass coverslips were washed with PBS and incubated with rabbit polyclonal anti-RFP antibody (a kind gift from Prof Ian Prior, University of Liverpool, 1:500 dilution in 1% (w/v) BSA in PBS for 30 min). Following incubation with primary antibody, cells were thoroughly washed in PBS and further incubated for 30 min with goat anti-rabbit Alexa 405- conjugated antibody (1:500, Life Technologies) in 1% (w/v) BSA in PBS. After final PBS washes, cells were fixed in 2% (w/v) formaldehyde in PBS for 6 min and mounted onto glass slides using ProLong^TM^ Gold Antifade glycerol (ThermoFisher Scientific).

### Lucifer Yellow (LY) assay

Cells were incubated in culture medium supplemented with 1 mg/ml LY carbohydrazide (CH) dipotassium salt (Santa Cruz) for 4 hours or overnight, at 37°C, 95% (v/v) air/5% (v/v) CO_2_, washed 3 times in PBS and chased in fresh culture medium for a minimum of 2 hours at 37°C, 5% (v/v) CO_2_. Cells were subsequently lysed and the incubation buffer collected as described in *Cell lysis and media collection for biochemical assays*. LY fluorescence was measured in the incubation buffer/cell lysate with excitation at 428 nm/emission at 536 nm wavelengths using a Jasco FP-6300 spectrofluorimeter.

### Cell lysis and media collection for biochemical assays

Cell incubation media were collected and stored on ice. All reagents and cells were maintained at 4°C throughout the lysis process. Cells were washed twice in PBS and incubated on ice on a rocking platform with RIPA buffer (ThermoScientifc^TM^) for 5 min. They were subsequently scraped from culture dishes and an equal volume of ice cold PBS added. Cellular debris were pelleted by centrifugation for 15 min at 13,000 rpm and the clarified lysates transferred to fresh tubes.

### N-Acetyl-β-D-Glucosaminidase (NAG) assay

Cells were subsequently lysed and the incubation buffer collected as described in *Cell lysis and media collection for biochemical assays*. Collected incubation buffers/cell lysates were incubated for 15 minutes to 1 hour at 37°C, 95% v/v air/5% v/v CO2 with 3 mM 4-nitrophenyl N-acetyl-β-D-glucosaminide, a NAG substrate, dissolved in 0.09 M citrate buffer solution (pH 4.5). The pH of the solution was systematically verified using Fisherbrand pH indicator paper sticks prior to NAG reaction. The reaction was stopped by adding 0.4 M sodium carbonate (pH 11.5). The product of the NAG substrate hydrolysis, *p*-nitrophenol, ionised upon addition of stop solution, was measured colorimetrically at 405 nm using a FLUOstar Omega Microplate Reader (BMG LABTECH).

### Cell transfection

All transfections were performed using Promofectin reagent (PromoKine) according to manufacturer’s instructions. Briefly, cells were cultured on 35 mm diameter MatTek glass bottom microscopy dishes or 13 mm diameter coverslips until they reached 50% confluence. Plasmid DNA (3 μg/MatTek dish, 1 μg/13 mm diameter coverslip) was mixed with 6 μl (MatTek) or 2 μl (coverslip) of Promofectin in serum-free RPMI (50 μl/μg DNA). The DNA/Promofectin solution was incubated for 20 minutes at room temperature then added dropwise to cells. Cells were assayed 24-48 hrs post-transfection.

### Imaging

Confocal imaging of fixed cells was performed on a Zeiss LSM 800 Airyscan microscope equipped with a Zeiss AxioObserverZ1, a 63x/1.4 Plan-Apochromat oil immersion objective and diode lasers as excitation light source (488 nm for pHluorin and Alexa 488, 405 for DAPI and Alexa 405 and 561 nm for mCherry). Emitted light was collected through Variable Secondary Dichroics (VSDs) onto a GaAsP-PMT detector.

TIRF imaging was performed on a 3i Marianas spinning-disk microscope equipped with a 100x/1.46 Alpha Plan-Apochromat oil objective, a Zeiss AxioObserver Z1, a 3i laserstack as excitation light source (488 nm for pHluorin and 561 nm for mCherry and Lysotracker®-red) and a TIRFM module. Emitted light was collected onto a CMOS camera (Hamamatsu, ORCA Flash 4.0) through quadruple bandpass. Live cells were maintained at 37 °C, 5 % CO_2_ (OKO lab incubation chamber).

Slidebook and Zen software suites were used for image acquisition. Images were processed and analysed using Fiji.

### Statistical analysis

Unless stated otherwise experiments were performed at least in triplicate (N = 3) and results are expressed as mean ± S.E.M. When applicable, the number of cells (n) is stated in the figure legends. All statistical analyses were performed on GraphPad Prism 6 software. Students’ unpaired *t*-tests was used for statistical comparison between groups, as indicated and p values in the figures are represented by stars (*p < 0.05, **p < 0.01 and ***p < 0.001).

## Acknowledgements

This work was supported by Wellcome Trust Prize PhD studentships awarded to CN and DR. All TIRFM and confocal microscopy studies were performed at The Biomedical Imaging Facility, Institute of Translational Medicine, University of Liverpool.

## Author contributions

CN performed experimental work, analysed the data and wrote the manuscript; NH wrote the manuscript and analysed the data; DR performed experimental work; RDB designed the study and wrote the manuscript; LPH designed the study, analysed the data and wrote the manuscript.

## Conflict of interest

The authors declare that they have no conflict of interest.

## Extended view figure legends

**Figure S1.**
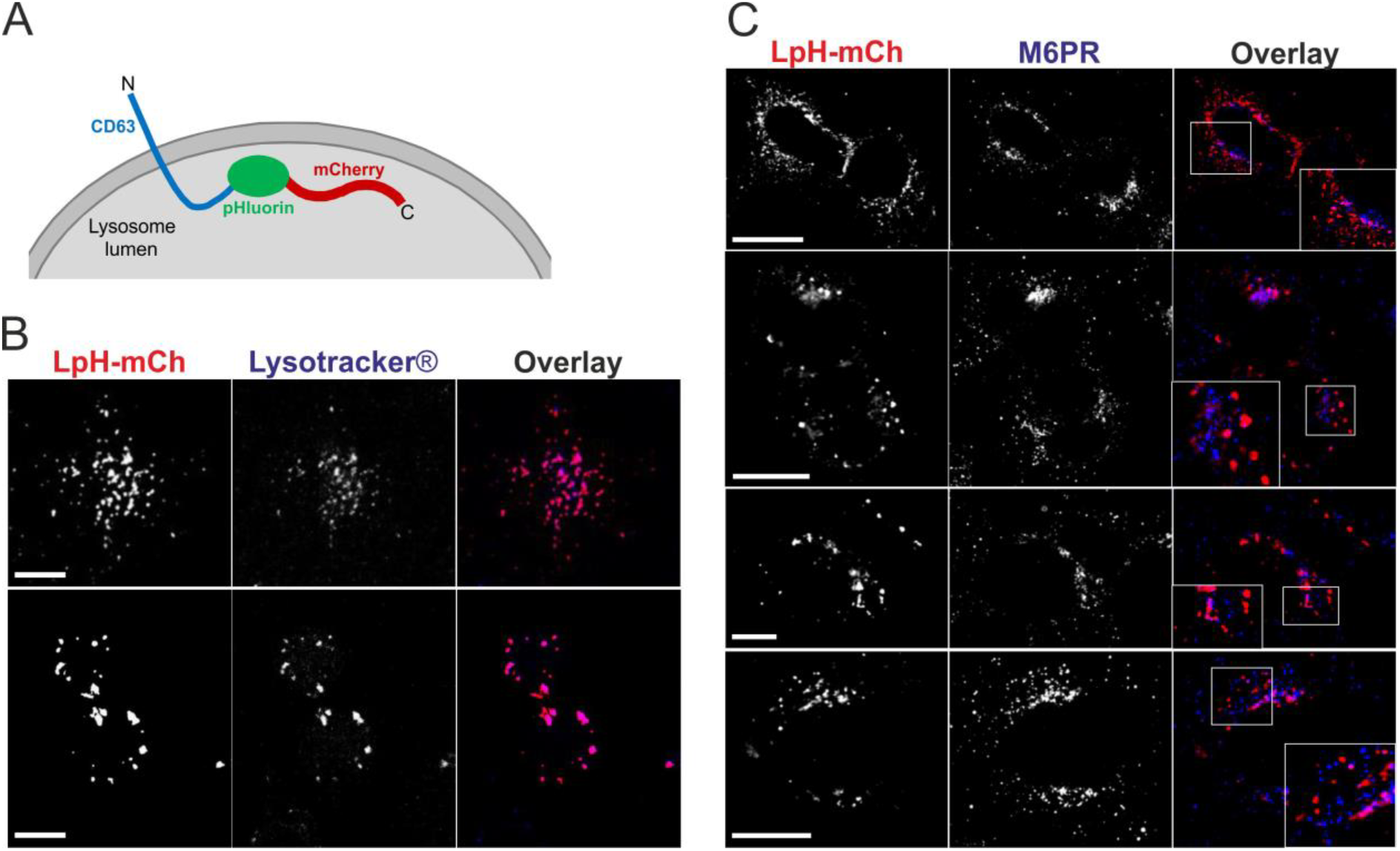
Architecture and cellular localisation of LysopHluorin-mCherry (LpH-mCh) in mammalian cells. A) Schematic representation of the LpH-mCh expression construct. CD63 targeting motif for localisation to lysosomes; pHluorin, a pH sensitive GFP-variant, quenched at acidic pH. The luminal mCherry tag, fluorescent at acidic pH, allows visualisation of the construct at all times. B) LpH-mCh (red) colocalises extensively with cellular organelles that accumulate the acidotropic dye Lysotracker® (blue). Regions of colocalisation appear pink in overlay images. C) LpH-mCh (red) exhibits limited colocalisation with endogenous mannose-6-phosphate receptor (α-M6PR antibody, blue). Four representative examples are shown. Scale bars 10 μm.

**Figure S2.**
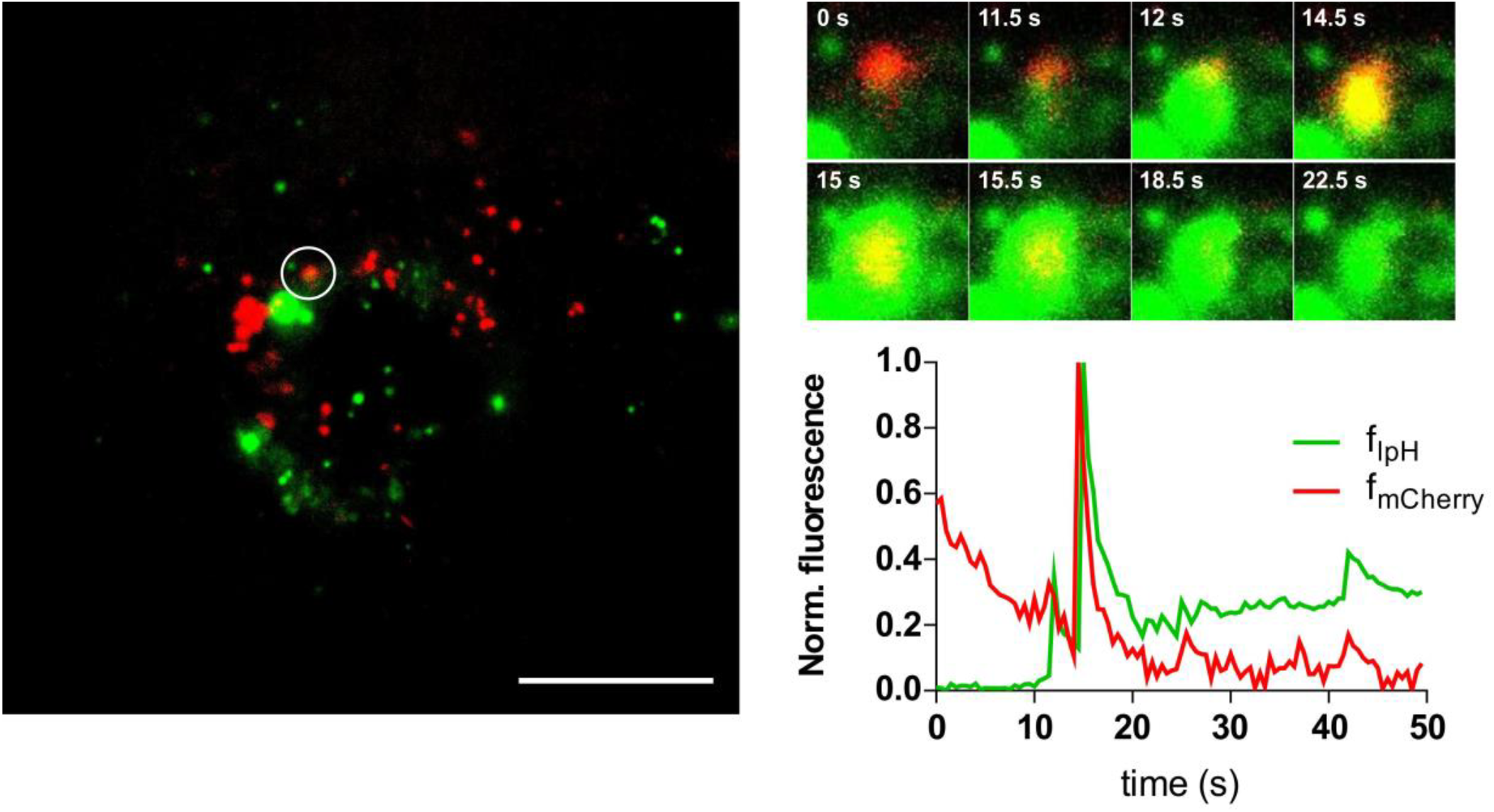
Characterisation of lysosome exocytosis in LpH-mCherry stable NRK cells. A) Representative TIRF data from a cell following treatment with 10 μM ionomycin and imaged immediately at a rate of 1 frame every 500 ms. A region of interest at the site of lysosome exocytosis was selected (white circle and expanded region of interest in (B)) and the fluorescence intensity of mCherry (fmCherry, red) and pHluorin (fLpH, green) measured. B) Exocytosis of lysosomes expressing LpH-mCherry at the PM was characterised by a de-quenching of pHluorin fluorescence which undergoes a sharp increase in intensity upon fusion, coincident with an increase in mCherry fluorescence (graph of normalised fluorescence intensity versus time and picture at frame t = 14.5 sec where a yellow signal is visible due to fluorescence in both mCherry and pHluorin channels). Diffusion of lysosomal membrane content in the plane of the PM is rapid and the yellow fluorescence signal is lost within 4s of exocytosis (t = 18.5 sec). The time on the zoomed pictures indicates the elapsed imaging time in sec. Scale bar 10 μm.

**Figure S3.**
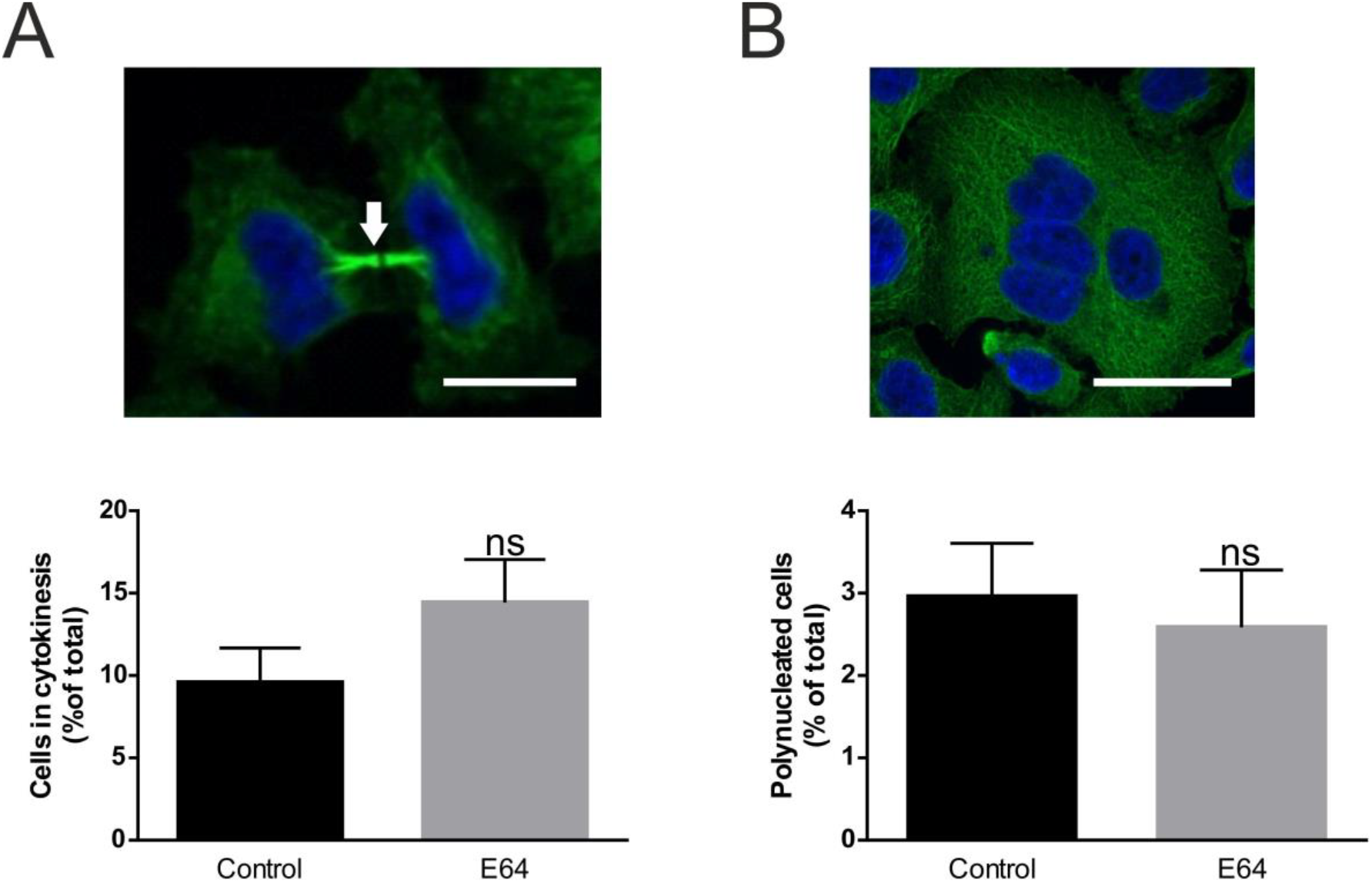
Inhibition of lysosomal cathepsins by E64 has no effect on HeLa cell mitosis. Cells were treated overnight with 100 μM E64 (or untreated, control), fixed and permeabilised. The proportion of cells linked by an intercellular bridge during cytokinesis (A) and the number of bi/tetranucleate cells (B) were assessed by spinning disk confocal microscopy using an anti-tubulin antibody (green) and DAPI (blue). n(control) = 960 cells, n(E64) = 909 cells, N = 6. The arrow indicates the position of the intercellular bridge. Unpaired student t-tests; results shown as ± S.E.M, ns: not significant. Scale bars in representative confocal sections: 10 μm.

**Movie S1**. *Lysosomes located near the plasma membrane localise to the site of cleavage furrow and cluster at either side of the cytoplasmic bridge.* BSC-1 cells were loaded with 50 nM Lysotracker®-red for 30 minutes and imaged on a TIRFM system (3i Marianas) at a rate of 1 frame every 30 seconds.

**Movie S2.** *Lysosomes cluster at either site of the intercellular cytoplasmic bridge during cytokinesis.* Examples of LpH-mCh-stable NRK cell undergoing cytokinesis. LpH-mCh-NRK cells were imaged every 0.5 s on a TIRFM system (3i Marianas).

**Movie S3.** *Lysosomes located near the plasma membrane localise to the site of cytoplasmic constriction and cluster at either side of the cytoplasmic bridge.* NRK cells were loaded with 50 nM Lysotracker®-red for 30 minutes and imaged on a TIRFM system (3i Marianas) at a rate of 1 frame every 2 seconds.

**Movies S4 and S5**. *Examples of lysosome exocytosis in LpH-mCh-stable NRK cells undergoing cytokinesis.* LpH-mCh stable NRK cells were imaged every 0.5 s on a TIRFM system (3i Marianas).

**Movie S6**. *Lysosome dynamics at interphase.* NRK cell was loaded with 50 nM Lysotracker®-red for 30 minutes and imaged on a TIRFM system (3i Marianas) at a rate of 1 frame every 2s.

**Movie S7**. *Characterisation of lysosome exocytosis at interphase.* LpH-mCh stably expressing NRK cells were treated with 10 μM ionomycin and imaged immediately on a TIRFM system (3i Marianas) at a rate of 1 frame every 0.5 s.

